# HUMAN ADENOVIRUS TYPE 4 COMPRISES TWO MAJOR PHYLOGROUPS WITH DISTINCT REPLICATIVE FITNESS AND VIRULENCE PHENOTYPES

**DOI:** 10.1101/2020.12.18.365858

**Authors:** Camden R. Bair, Wei Zhang, Gabriel Gonzalez, Arash Kamali, Daniel Stylos, Jorge C. G. Blanco, Adriana E. Kajon

## Abstract

Human adenovirus type 4 (HAdV-E4) is the only type (and serotype) classified within species *Human mastadenovirus E* that has been isolated from a human host to the present. Recent phylogenetic analysis of whole genome sequences of strains representing the spectrum of intratypic genetic diversity described to date identified two major evolutionary lineages designated phylogroups (PG) I, and II, and validated the early clustering of HAdV-E4 genomic variants into two major groups by low resolution restriction fragment length polymorphism analysis. In this study we expanded our original analysis of intra- and inter-PG genetic variability, and used a panel of viruses representative of the spectrum of genetic diversity described for HAdV-E4 to examine the magnitude of inter- and intra-PG phenotypic diversity using an array of cell-based assays and a cotton rat model of HAdV respiratory infection. Our proteotyping of HAdV-E strains using concatenated protein sequences in selected coding regions including E1A, E1B-19K and −55K, DNA polymerase, L4-100K, various E3 proteins, and E4-34K confirmed that the two clades encode distinct variants/proteotypes at most of these loci. Our *in vitro* and *in vivo* studies demonstrated that PG I and PG II differ in their growth, spread, and cell killing phenotypes in cell culture and in their pulmonary pathogenic phenotypes. Surprisingly, the differences in replicative fitness documented *in vitro* between PGs did not correlate with the differences in virulence observed in the cotton rat model. This body of work is the first reporting phenotypic correlates of naturally occurring intratypic genetic variability for HAdV-E4.

**IMPORTANCE:** Human adenovirus type 4 (HAdV-E4) is a prevalent causative agent of acute respiratory illness of variable severity and of conjunctivitis and comprises two major phylogroups that carry distinct coding variations in proteins involved in viral replication and modulation of host responses to infection. Our data show that PG I and PG II are intrinsically different regarding their ability to grow and spread in culture and to cause pulmonary disease in cotton rats.

This is the first report of phenotypic divergence among naturally occurring known genetic variants of a HAdV type of medical importance. This research reveals readily detectable phenotypic differences between strains representing phylogroups I and II, and it introduces a unique experimental system for the elucidation of the genetic basis of adenovirus fitness and virulence and thus for increasing our understanding of the implications of intratypic genetic diversity in the presentation and course of HAdV-E4-associated disease.

## INTRODUCTION

Human adenovirus type 4 (HAdV-E4), the only type (and serotype) classified within species *Human mastadenovirus E* (HAdV-E) isolated from humans to the present, was originally recovered from a respiratory specimen obtained from a military recruit in basic training in the winter of 1952-1953 in Fort Leonard Wood, MO, USA during an epidemic outbreak of acute respiratory disease (ARD) (1). The “new” virus was designated RI-67, as it was derived from Respiratory Illness case number 67 in the epidemic (1). The circulation of HAdV-E4 was subsequently documented between 1958 and 1959 in Taiwan in association with cases of conjunctivitis and ARD (2). By the 1980s this unique type had been widely recognized as a prevalent causative agent of respiratory illness of variable severity, and also of conjunctivitis with clinical presentations ranging from mild “pink eye” to severe epidemic keratoconjunctivitits (EKC) (3–11).

The pioneer investigations of intratypic genetic variability for HAdV-E4 conducted in the late 1980s - early 1990s using low resolution restriction fragment length polymorphism (RFLP) analysis demonstrated the existence of two major clusters of genomic variants readily distinguishable by their digestion profiles with BamHI, DraI, EcoRV, and SmaI, among other endonucleases (12, 13). Based on their BamHI digestion profiles, these variants were designated by Li and Wadell (12) with a letter “p” for prototype-like, or a letter “a” or “b” for the non-prototype like variants followed by numbers 1, 2, etc., designating additional variability detectable with other endonucleases. Our recent evaluation of a sizable collection of whole genome sequences (n=47) representing 45 strains isolated worldwide from clinical specimens between 1953 and 2015 (14) and the spectrum of genetic variability described to date by RFLP (11–13, 15) provided a global view of the magnitude of genetic diversity for this medically important type, confirmed the early groupings of related genomic variants, and identified two major evolutionary lineages designated phylogroups (PG) I, and II (14). PG I and PG II comprise the prototype-like and non-prototype-like genomic variants, respectively.

The two PGs differ significantly in their G + C content and apparent evolutionary rates, and as we recently reported, they also exhibit marked sequence differences in coding regions including but not limited to E1A, E1B, E3, L3, and L4 (14).

Interestingly, published data indicate that the representation of PG II strains among collections of respiratory and ocular clinical specimens sampled from symptomatic individuals in a variety of settings since 1965 has greatly exceeded that of PG I strains (11, 15–18) suggesting a higher fitness and/or possibly a higher virulence and/or transmissibility for these viruses.

To the present, no studies have formally identified genetic determinants of adenovirus virulence and fitness. Importantly, the pathobiological implications for the intratypic genetic variability documented for HAdV-E4 and other types remain unknown.

In this study, we expand our original analysis of intra- and inter-PG genetic variability (14) and use a panel of viruses representative of the spectrum of genetic diversity described for HAdV-E4 to examine the magnitude of inter- and intra-PG phenotypic diversity using an array of cell-based assays and a cotton rat model of HAdV respiratory infection.

Our data show that PG I and PG II display distinct coding variations in proteins involved in viral replication and modulation of host responses to infection, and that they are intrinsically different regarding their ability to grow and spread in culture and to cause pulmonary disease in cotton rats.

## RESULTS

### Deeper analyses of genomic sequences for HAdV-E4 strains in PG I and PG II reveal additional genetic differences between and within the two major clades

The use of new analytical tools to further assess variation across the 45-genome collection originally characterized by Gonzalez *et al*. (14) revealed genome-wide sequence differences between strains clustering in PG I and PG II and significant diversity in protein-coding sequences (Fig 1A). A computational analysis of the encoded proteome (proteotyping) of these 45 HAdV-E strains using concatenated predicted protein sequences in selected coding regions including E1A, E1B 19K and 55K, DNA polymerase, L4 100K, E3 proteins CR1 α, gp19K, CR1β, CR1 δ, RIDα, RIDβ and 14.7K, and E4 34K (Fig 1B) confirmed that the two clades encode distinct variants/proteotypes at most of these loci. Importantly, in each phylogroup additional divergence was identified with the occurrence of minor proteotypes; for example, strains USA 2000 (AP014852), USA 2004 (AP014853) and USA 2011 (EF371058) of PG-I show characteristic mutations in E2B DNA polymerase, L4 100K, and proteins in the E3 region, reflecting the emergence and maintained circulation in the USA of a distinct lineage. Intra-phylogroup II variations were also identified, with multiple small groups of sequences showing characteristic mutations across the DNA polymerase and E3 proteins.

**Figure 1:**
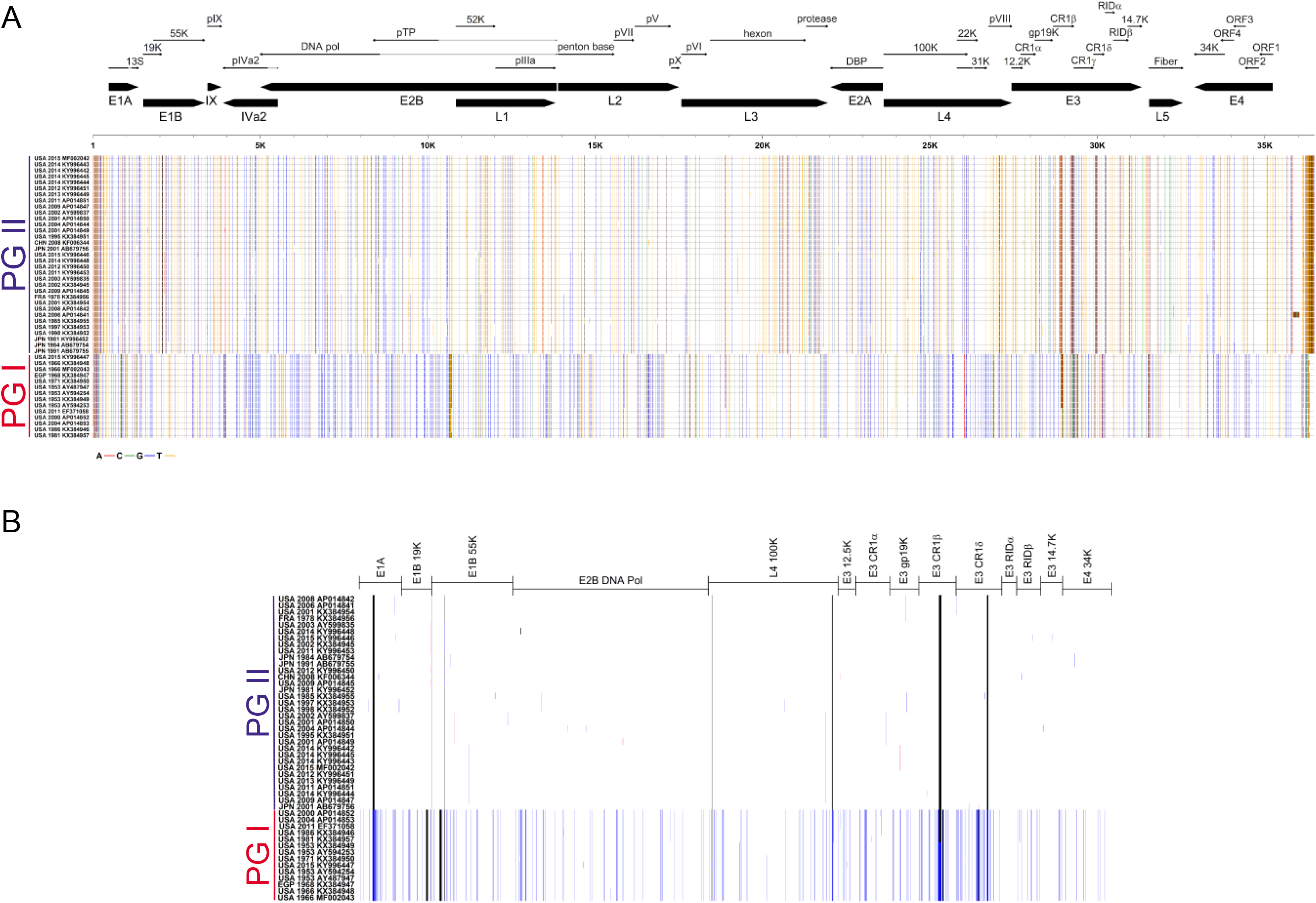
**A. Sequence differences among HAdV-E4 strains of phylogroups I and II (PG I and PG II).** The consensus was obtained from the multiple sequence alignment built using MAFFT (63) with the set of whole genome sequences originally analyzed by Gonzalez *et al*. (15), and summarized with the R package APE to identify by position the consensus nucleotide and the mutations. The GenBank accession numbers for the examined genomic sequences together with the place and year of isolation of the corresponding strain are shown on the vertical axis on the left. Positions with nucleotide mutations differing from the consensus are colored with red, green, blue or orange, corresponding to mutations to adenine, cytosine, guanine or thymine, respectively. **B. Proteotyping of concatenated predicted protein sequences for selected coding regions.** The GenBank accession numbers for the examined genomic sequences together with the place and year of isolation of the corresponding strain are shown on the vertical axis on the left. The start, end and polypeptide name of the concatenated sequences arranged in genome order are shown on the top of the panel. Sites with amino acid polymorphisms are colored according to the frequency across sequences, with blank for the consensus, blue and red for the first and second most frequent polymorphisms for a position; gap sites are colored in black.

### PG I and PG II strains differ in their average plaque size and cell-to-cell spread phenotype

A panel of HAdV-E4 strains representative of the spectrum of HAdV-E4 genomic variants were selected for comparison of *in vitro* growth characteristics. Proteotyping results for coding regions in genome order for this subset of strains are shown in Table 1.

**Table 1:** Genome-wide proteotyping results for the seven HAdV-E4 strains examined in this study

|  |  |  |  | PG I |  |  |  | PG II |  |  |
| --- | --- | --- | --- | --- | --- | --- | --- | --- | --- | --- |
|  |  |  |  | RI-67<sup>a</sup> | CL68578 | NY-15-4054 | NHRC 90339 | NHRC 22652 | NY-12-27440 | NHRC 42606 |
| Gene | Product | Amino acid diversity | No. of variants | AY594253<sup>b</sup> | AY594254 | KY996447 | EF371058 | AP014843 | KY996451 | AY599835 |
| E1A | 13S | 0.04 ± 0.04 | 2 | v1<sup>f</sup> | v1 | v1 | v1 | v2 | v2 | v2 |
| E1B | 19K | 0.03 ± 0.03 | 2 | v1 | v1 | v1 | v1 | v2 | v2 | v2 |
|  | 55K | 0.01 ± 0.01 | 3 | v1 | v1 | v1 | v1 | v2 | v2 | v2 |
| IX | pIX | 0.01 ± 0.01 | 2 | v1 | v1 | v1 | v1 | v2 | v2 | v2 |
| IVa2 | IVa2 | 0.01 ± 0.01 | 2 | v1 | v1 | v1 | v1 | v2 | v2 | v2 |
| E2B | DNA pol<sup>c</sup> | 0.01 ± 0.02 | 3 | v1 | v1 | v2 | v2 | v3 | v3 | v3 |
|  | pTP<sup>d</sup> | 0.01 ± 0.01 | 2 | v1 | v1 | v1 | v1 | v2 | v2 | v2 |
| L1 | 52-55KDa | 0.01 ± 0.01 | 2 | v1 | v1 | v1 | v1 | v2 | v2 | v2 |
|  | pIIIa | 0.01 ± 0.01 | 2 | v1 | v1 | v1 | v1 | v2 | v2 | v2 |
| L2 | pIII | 0.01 ± 0.01 | 3 | v1 | v1 | v1 | v1 | v2 | v2 | v2 |
|  | pV | 0.02 ± 0.02 | 2 | v1 | v1 | v1 | v1 | v2 | v2 | v2 |
|  | pVII | 0.01 ± 0.01 | 2 | v1 | v1 | v1 | v1 | v2 | v2 | v2 |
|  | pX | 0 ± 0 | 1 | v1 | v1 | v1 | v1 | v1 | v1 | v1 |
| L3 | pVI | 0.01 ± 0.01 | 2 | v1 | v1 | v1 | v1 | v2 | v2 | v2 |
|  | hexon | 0.01 ± 0.01 | 3 | v1 | v1 | v3 | v1 | v2 | v2 | v2 |
|  | protease | 0.02 ± 0.02 | 2 | v1 | v1 | v1 | v1 | v2 | v2 | v2 |
| E2A | DBP<sup>e</sup> | 0.01 ± 0.01 | 2 | v1 | v1 | v1 | v1 | v2 | v2 | v2 |
| L4 | 100KDa | 0.01 ± 0.02 | 6 | v1 | v1 | v1 | v2 | v3 | v3 | v3 |
|  | 33KDa | 0.01 ± 0.01 | 2 | v1 | v1 | v1 | v1 | v2 | v2 | v2 |
|  | pVIII | 0.01 ± 0.01 | 2 | v1 | v1 | v1 | v1 | v2 | v2 | v2 |
| E3 | 12.1KDa | 0.02 ± 0.03 | 3 | v1 | v1 | v2 | v2 | v3 | v3 | v3 |
| | CR1α | 0.02 ± 0.03 | 2 | v1 | v1 | v1 | v1 | v2 | v2 | v2 |
|  | gp19K | 0.02 ± 0.02 | 3 | v1 | v1 | v1 | v1 | v2 | v2 | v2 |
| | CR1β | 0.04 ± 0.05 | 3 | v1 | v1 | v1 | v2 | v3 | v3 | v3 |
| | CR1δ | 0.04 ± 0.05 | 3 | v1 | v1 | v1 | v1 | v2 | v2 | v2 |
| | RIDα | 0 ± 0 | 1 | v1 | v1 | v1 | v1 | v1 | v1 | v1 |
| | RIDβ | 0.03 ± 0.03 | 3 | v1 | v1 | v1 | v2 | v3 | v3 | v3 |
|  | 14.7K | 0.01 ± 0.01 | 2 | v1 | v1 | v1 | v1 | v2 | v2 | v2 |
| L5 | fiber | 0.01 ± 0.01 | 2 | v1 | v1 | v1 | v1 | v2 | v2 | v2 |
| E4 | 34K | 0.01 ± 0.01 | 2 | v1 | v1 | v1 | v1 | v2 | v2 | v2 |
|  | 13.5K | 0.03 ± 0.03 | 2 | v1 | v1 | v1 | v1 | v2 | v2 | v2 |
|  | 13.7K | 0 ± 0 | 2 | v1 | v1 | v1 | v1 | v1 | v1 | v1 |
|  | 14.1K | 0.01 ± 0.02 | 2 | v1 | v1 | v1 | v1 | v2 | v2 | v2 |
|  | 14.6K | 0.03 ± 0.04 | 2 | v1 | v1 | v1 | v1 | v2 | v2 | v2 |
|  | 7.4K | 0.03 ± 0.04 | 2 | v1 | v1 | v1 | v1 | v2 | v2 | v2 |

a. Strain designation; b. GenBank accession number for whole genome sequence; c. DNA polymerase; d. precursor of the terminal protein; e. DNA binding protein; f. protein variant/ 1, 2, 3. variant number

First, we assessed whether the stocks for the selected 7 strains exhibited any detectable differences in specific infectivity. The number of genome copies per PFU for each of the strains was determined by two independent quantitative PCR (qPCR) approaches: the DiaSorin qPCR platform described in the Methods section and the protocol developed by Lu *et al*. at the US Centers for Disease Control and Prevention (19). No significant differences in the number of genome copies per PFU were detectable between individual strains or PGs by either methodology (data not shown).

Next, we examined plaque phenotypes for the selected 7 HAdV-E4 strains in A549 human alveolar epithelial A549 cells. As shown in Figure 2, no obvious differences in plaque morphology were detected by visual examination of bright field images taken at 7 days post infection (dpi). However, PG I strains had larger plaque sizes than PG II strains. The vaccine strain CL68578 formed the largest plaques, while strains NY-15-4054, the cell culture-adapted RI-67 strain, and NHRC 90339 formed relatively smaller plaques. The PG II strains formed uniformly smaller plaque sizes that were not significantly smaller than those formed by the PG I strain NHRC 90339. One-way ANOVA with Holm-Sidak’s multiple comparisons test showed significant differences between PG I strains CL68578, NY-15-4054, and RI-67, and all 3 examined PG II strains as well as PG I strain NHRC 90339.

**Figure 2:**
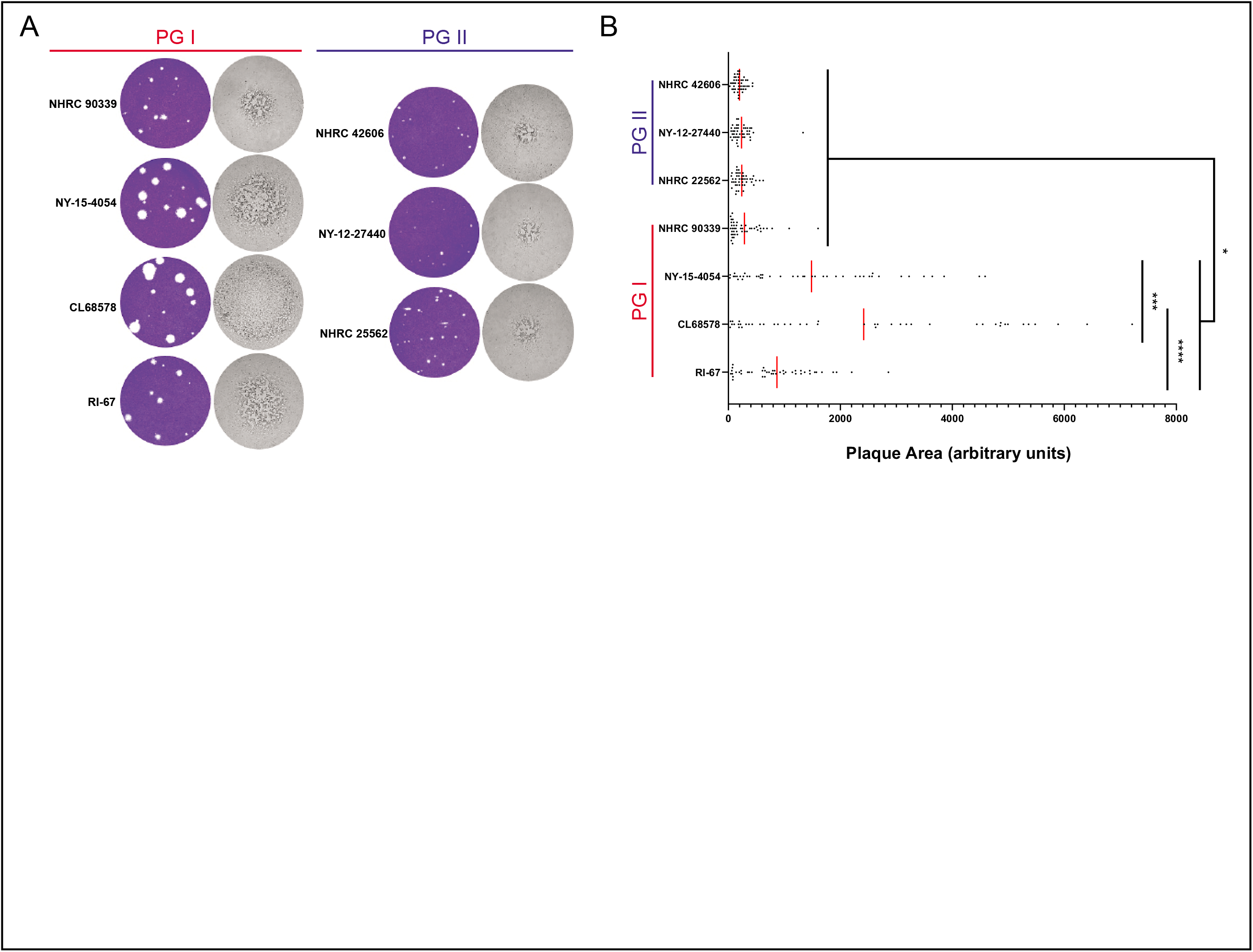
Plaque phenotypes. (A) Plaque assays on A549 cell monolayers in 6-well plates. Monolayers infected with PG I (RI-67, CL68578, NY-15-4054, and NHRC 90339) or PG II (NHRC 25562, NY-12-27440, and NHRC 42606) strains were fixed and stained with crystal Violet at 7 days post infection. For each examined strain, representative brightfield images of individual plaques are shown next to the corresponding images of a representative crystal-violet stained monolayer. (B) Plaque areas (in arbitrary units) were determined using FIJI image analysis software. Statistical significance was assessed by one-way ANOVA (6, 484 = 37.5) and Holm-Sidak’s multiple-comparison test (*, p < 0.05; ***, p < 0.001; ****, p < 0.0001).

The rate of plaque development was examined by calculating the percentage of plaques detectable on any given day post infection relative to the final total number of plaques over a period of 25 days, following the approach used by Tollefson *et al*. to characterize HAdV-C ADP mutants (20, 21). As shown in Fig 3A, plaques formed by PG I strains were visible one day earlier (day 4) than plaques formed by PG II strains (day 5). PG I strains CL68578 and NY-15-4054 exhibited the fastest rate of plaque development. At 10 dpi, about 90% and 70% of all detected plaques were visible for the CL68578 and NY-15-4054, respectively, while only 30-40% of all detected plaques were visible for the other examined strains, including NHRC 90339.

**Figure 3:**
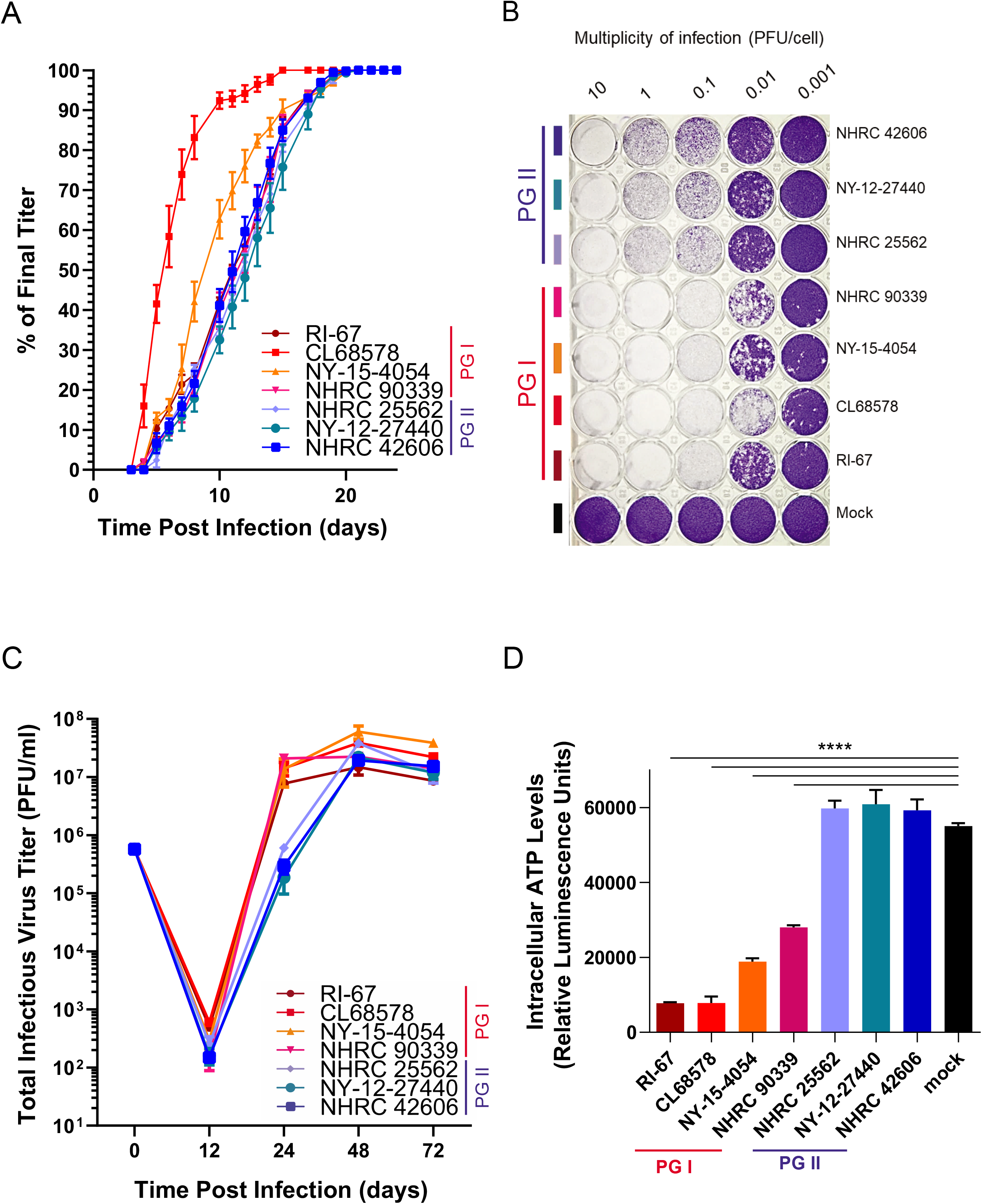
Growth and spread phenotypes. **A. Plaque development.** Plaques newly developed on infected A549 cell monolayers in 60 mm dishes were counted every one to two days. The total number of plaques observed on a given day is plotted as a percentage of the total number of plaques observed on the final day of the assay. **B. Lateral spread.** A549 cell monolayers were infected at MOIs ranging from 0.001 to 10 PFU/cell with the indicated PG I or PG II strains. At 7 days post infection, cell monolayers were fixed and stained with crystal violet for visualization of the extent of cytopathic effect. **C. Replication kinetics.** Submerged A549 cell monolayers were infected with PG I or PG II strains at a MOI of 1 PFU/cell. Infectious virus yields were determined at the indicated time points post infection by plaque assay. **D. Cell killing phenotype.** A549 cells were infected at a MOI of 10 PFU/cell. At 96 hours post infection, intracellular ATP levels were measured as indicators of cell viability. Error bars denote SEM. Statistical significance was accessed by one-way ANOVA analysis, F (7, 24) = 140.9, p < .0001. Tukey’s post hoc comparisons to mock-treated cells revealed a significant difference. ****, p < .0001.

To evaluate cell-to-cell spread phenotypes, cytopathic effect (CPE) progression was examined over a period of 7 days in A549 cell monolayers infected at different multiplicities (MOI 0.001 - 10 PFU/cell) as described by Doronin *et al*. (22, 23). Overall, the three PG II strains spread at a restricted rate compared to the PG I strains (Fig 3B).

### Differences in replication kinetics and cell killing account for the observed differences in plaque size and spread phenotypes

The larger plaque phenotype and enhanced spread phenotype of PG I strains could either be a reflection of a faster replication kinetics and/or of enhanced cytopathic effect resulting from infection. To investigate this, we compared the growth kinetics of PG I and PG II strains in A549 cell monolayers infected at a MOI of 1 PFU/cell. While infectious virus yields were similar at 48 and 72 hours post infection (hpi), infectious virus yields at 24 hpi were higher for PG I strains (Fig 3C), suggesting faster growth kinetics. Next, we examined cell killing phenotypes in A549 cell monolayers infected with PG I and PG II strains at a MOI of 10 PFU/cell. ATP levels were determined as indicators of cell viability. At 4 dpi intracellular ATP levels in PG II-infected cells were relatively high, and similar to those detected in mock-infected cells. In contrast, in PG I-infected cells ATP levels were significantly lower, reflecting increased cell death. ATP levels were the lowest in RI-67- and CL68578-infected cells and NY-15-4054- and NHRC 90339 exhibited intermediate cell killing phenotypes. (Fig 3D). From the results of these experiments we conclude that both faster growth kinetics and increased killing may contribute to the documented large plaque and lateral spread phenotypes of PG I strains.

### Examination of PG I and PG II strains in a cotton rat model of adenovirus respiratory infection revealed differences

For an initial comparison of pathogenic phenotypes of PG I and PG II *in vivo* we selected one strain from each PG to represent the most extreme plaque size, spread, and cell killing phenotypes observed: CL68578, (PG I) and NHRC 42606 (PG II). We first conducted some additional cell-based assays to expand the characterization of these two strains. As shown in Figure 4A, and consistent with our initial findings, PG I strain CL68578 exhibited a faster kinetics of viral progeny release than NHRC 42606 in polarized human airway epithelial Calu-3 cells. Moreover and consistent with the results of the cell viability assays based on intracellular ATP detection, significantly higher levels of lactate dehydrogenase (LDH) activity indicative of enhanced cell death (24) were detected at 48 and 72 hpi in supernatants of A549 cells infected with CL68578 (Fig 4B). Both strains replicated in cotton rat lung epithelial cells - albeit at modest levels compared to those observed in A549 cells, with CL68578 displaying faster growth kinetics (Fig 4C).

**Figure 4:**
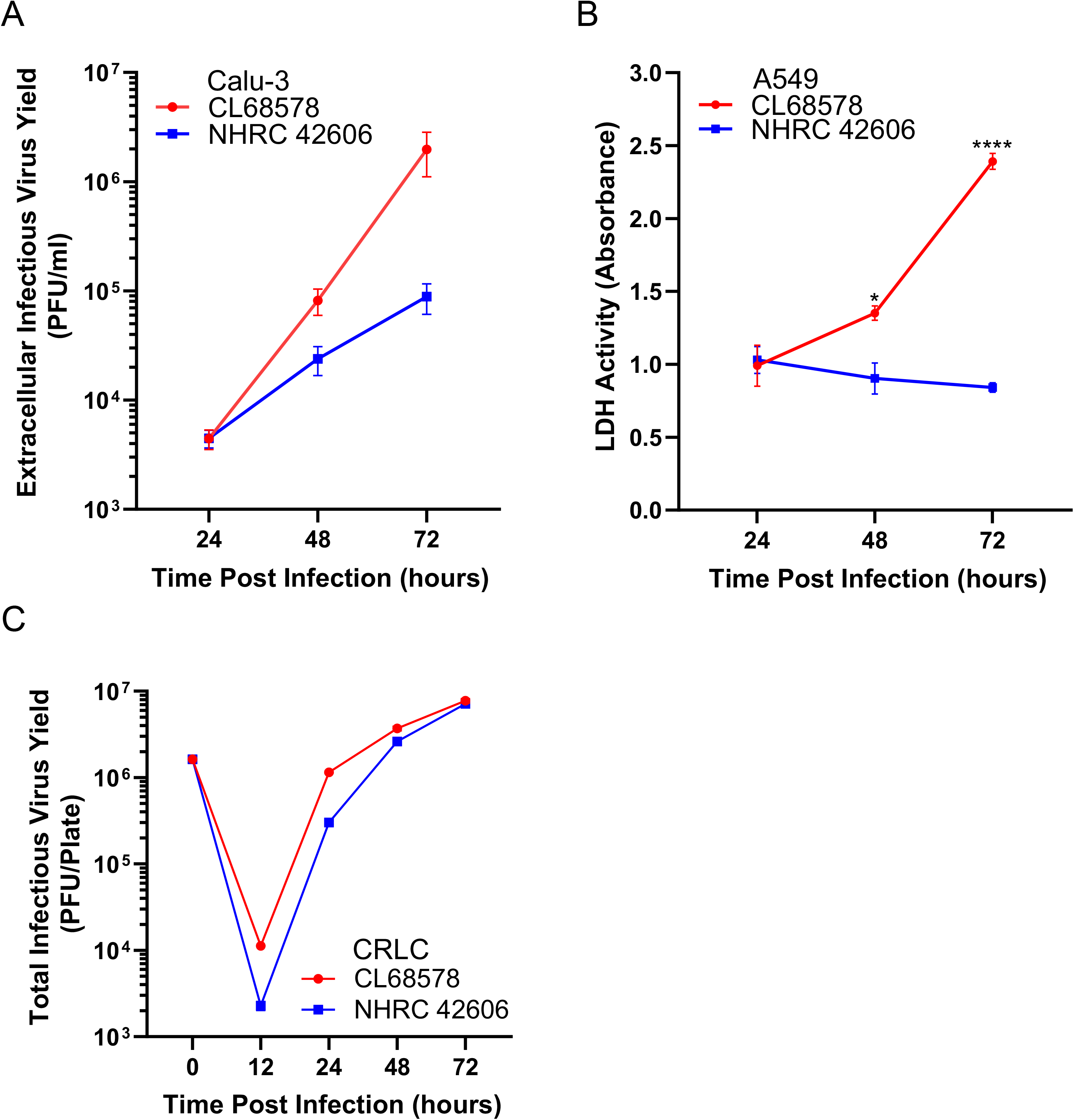
Additional phenotypic characterization of strains CL68578 and NHRC42606 using cell-based assays. **A. Kinetics of viral progeny release in polarized human airway epithelial cells.** Polarized Calu-3 cells in transwell inserts were infected at a MOI of 10 PFU/cell. At the indicated times post infection, the apical compartment was washed twice with a total of 400 μl of PBS and infectious virus titers in the combined wash fluid were determined by plaque assay. **B. Kinetics of LDH release from infected A549 cells.** A549 cell monolayers in 48-well plates were infected at a MOI of 1 PFU/cell in biological triplicates. LDH activity in infected cell supernatants was determined as an indicator of cell lysis. Statistical significance was assessed using the Student’s *t* test, *p< 0.05, **** p < 0.0001. **C. Growth kinetics in monolayers of cotton rat lung epithelial cells.** CRLC were infected at a MOI of 5 PFU/cell in biological triplicates. Infectious virus yields were determined by plaque assay in A549 cells.

We infected female cotton rats intranasally (i.n.) with 10^7^ PFU of strains CL68578 or NHRC 42606 (n=3 rats per group) and found marked differences in the magnitude of the resulting pulmonary pathology at day 3 pi (data not shown). In a second study, female cotton rats in groups of 6 were infected with 10^7^ PFU of each virus i.n. and euthanized at 1, 2, 3, and 6 dpi. The assessment of viral load in infected cotton rat lungs revealed differences in the course of infection between the two examined viruses. Two of the animals infected with NHRC 42606 (PG II) pre-segregated into two different cages died at 3 dpi. These animals could not be dissected for tissue sampling and were therefore excluded from subsequent analysis. Similar numbers of genome copies per gram of tissue were detected by real-time quantitative PCR (qPCR) at 1 dpi in the lungs of rats infected with the PG I or PG II strains (Fig. 5A). The viral load increased modestly by 2 dpi in animals infected with CL68578 (PG I) and gradually declined thereafter. Consistent with the *in vitro* growth phenotypes documented for PG II strain NHRC 42606, lung viral loads were almost a log unit lower at 2, 3, and 6 dpi in the animals infected with this virus.

**Figure 5:**
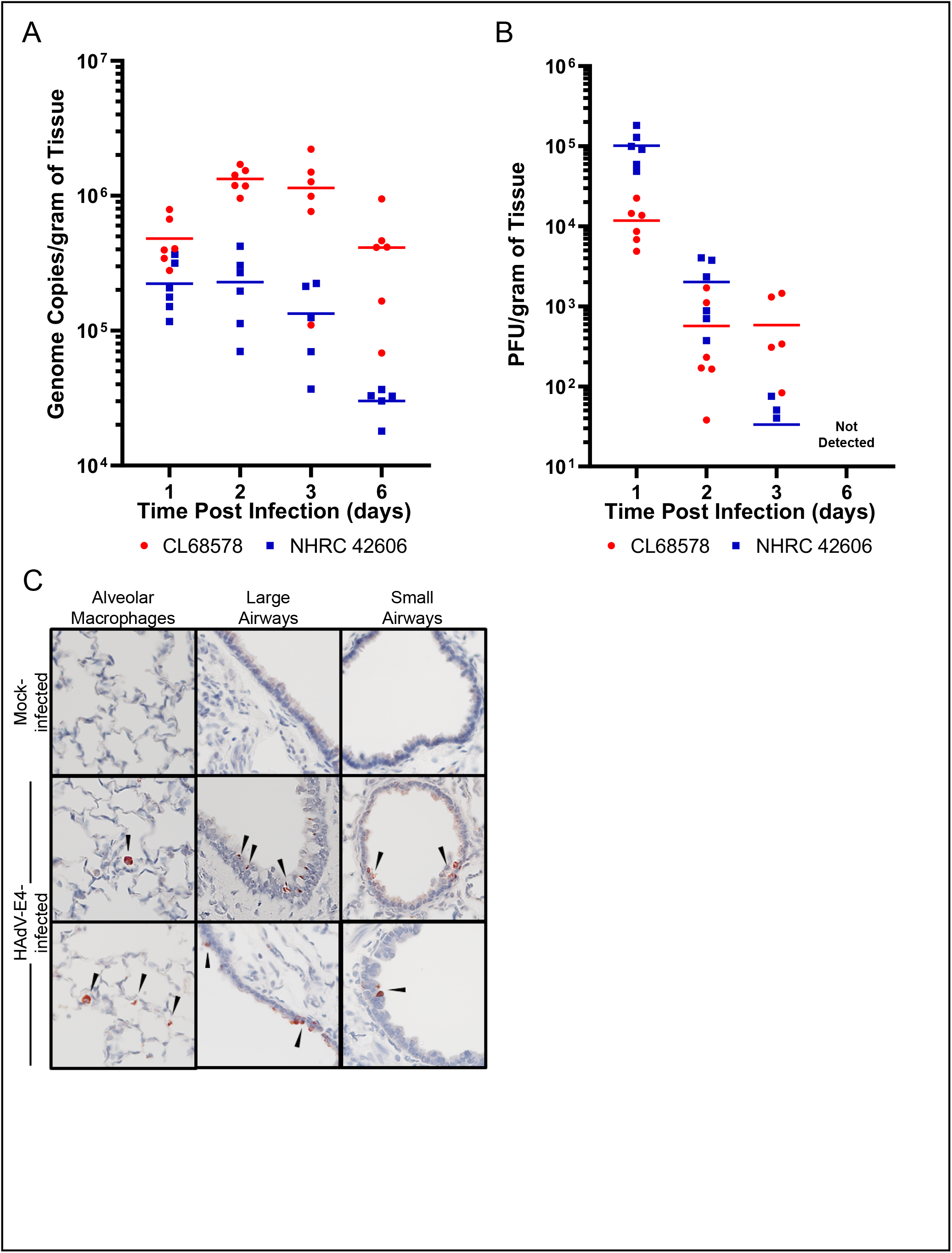
Progression and localization of infection in cotton rat lungs. **A. Assessment of viral load in the lung of infected cotton rats by qPCR.** The viral load in the supernatants of left lung lobe homogenates from PG I and PG II-infected cotton rats at the indicated times post infection was determined by qPCR using a DiaSorin Molecular proprietary platform for human adenovirus. Genome copy numbers were normalized to the mass of homogenized tissue. Horizontal bars denote mean values for each group. Each symbol represents a single animal. **B. Infectious virus load in the lungs of infected cotton rats.** Infectious virus loads in lung homogenate supernatants were determined by plaque assay. Titers were normalized to the mass of homogenized tissue and loads expressed as PFU/gram of tissue. Horizontal bars denote mean values for each group. Infectious virus was not detectable in samples collected at 6 dpi. **C. Immunohistochemical detection of hexon protein in lung sections of infected cotton rats.** Paraffin-embedded lung sections on glass slides were stained with AdV Mab MAB805 (Millipore). Images from stained sections were acquired in a Nikon Eclipse E600 microscope equipped with a Nikon DXM1200F digital camera at 40X magnification. Arrowheads indicate positive cells.

Infectious virus was detectable by plaque assay in lung homogenates of infected rats in both groups until day 3 pi in 4/6 animals infected with CL68578 and in 3/5 surviving animals infected with NHRC 42606 (Fig. 5B). While titers at 1 dpi were higher in rats infected with PG II strain NHRC 42606, a steady decline in viral load was documented for this group supporting the qPCR results and suggesting a more rapid clearance of the virus. HAdV hexon protein was readily detectable by immunohistochemistry in very modest amounts in large and small airway epithelial cells and alveolar macrophages during the first 3 days after infection (Fig. 5C) confirming that the cotton rat is a permissive host for HAdV-E4. This assay revealed another difference between the examined PG I and PG II strains, the localization of the signal in infected animals, particularly at 1 dpi (Table 2). At this time, while the virus was almost exclusively detected in airway epithelia in the rats infected with the PG I strain, the signal was predominantly found in alveolar macrophages in the rats infected with the PG II strain. AdV hexon protein was also detectable in alveolar macrophages of rats infected with the PG II strain on days 2 and 3 post infection.

**Table 2:** Localization of viral infection in cotton rat lungs during the first 3 days post infection^a^

| virus | dpi<sup>b</sup> | Detection of HAdV hexon protein by immunohistochemistry: |  |  |
| --- | --- | --- | --- | --- |
|  |  | Alveolar macrophages | Large airways | Small airways |
| vehicle | 1-6 | 0/6 rats | 0/6 rats | 0/6 rats |
| PG I<br>CL68578 | 1 | 1/6 rats | 6/6 rats | 6/6 rats |
|  | 2 | 0/6 rats | 6/6 rats | 6/6 rats |
|  | 3 | 0/6 rats | 6/6 rats | 6/6 rats |
| PG II<br>NHRC 42606 | 1 | 6/6 rats | 6/6 rats | 6/6 rats |
|  | 2 | 6/6 rats<sup>c</sup> | 6/6 rats | 6/6 rats |
|  | 3<sup>d</sup> | 2/5 rats | 5/5 rats | 4/5 rats |

To assess the extent of pulmonary inflammation resulting from HAdV-E4 respiratory infection, lung sections from mock- and HAdV-E4-infected animals were stained with hematoxylin and eosin, and scored for several metrics of pulmonary pathology as described in Materials and Methods. Representative photomicrographs of stained lung sections from rats euthanized at 1, 2, 3, and 6 dpi are shown in Figure 6A.

**Figure 6:**
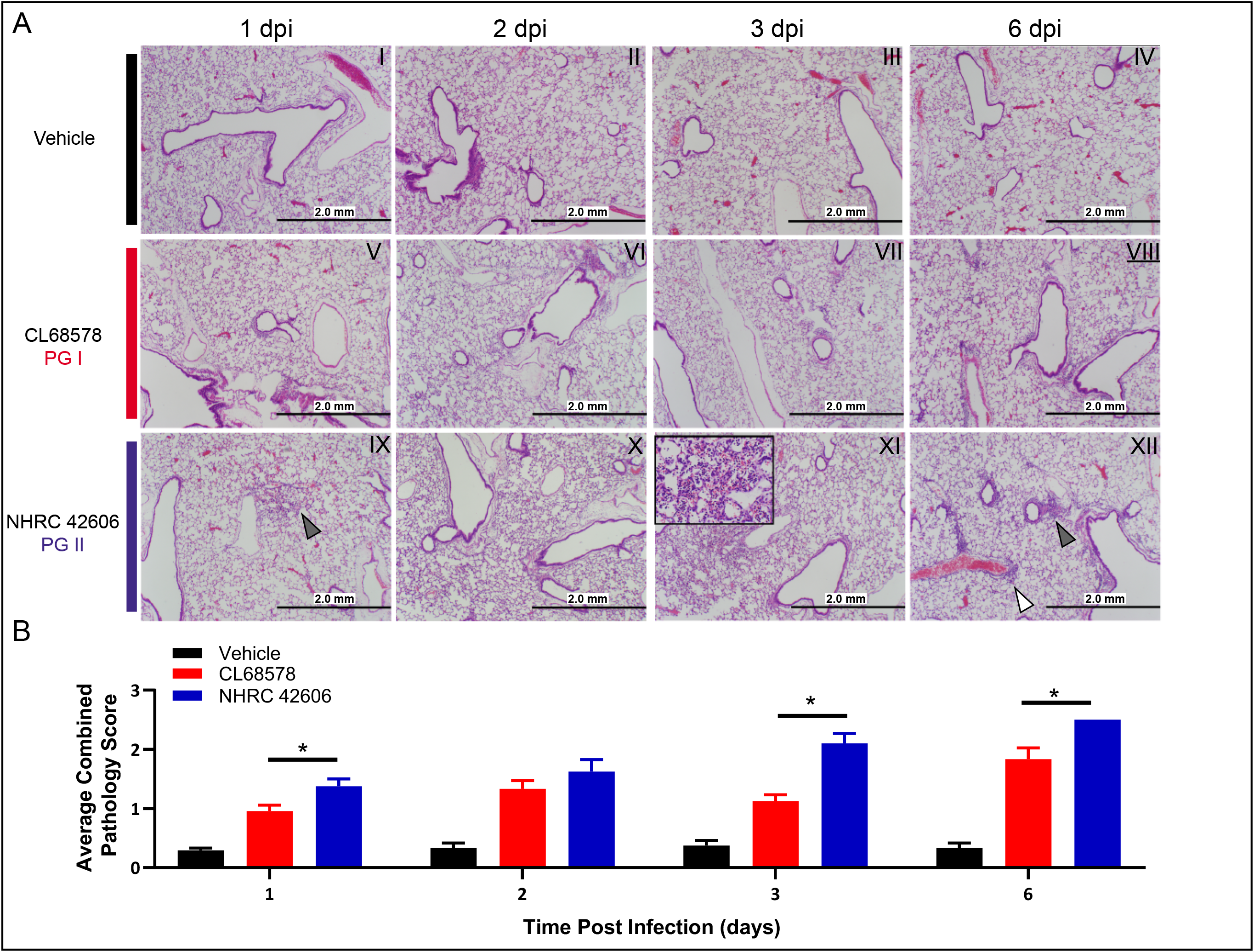
Pulmonary pathology in cotton rats infected intranasally. **A. Microscopic examination of lung sections.** The right lung lobes of mock- or HAdV-E4-infected cotton rats were harvested at 1, 2, 3 and 6 dpi and processed for paraffin-embedding, sectioning and hematoxylin and eosin staining for microscopic comparative examination of lung pathology. Representative photomicrographs for each experimental group and time post infection were taken at a 40X magnification in an Olympus BH-2 microscope equipped with an Olympus DP70 digital camera and Olympus DP Controller software with focus on terminal bronchioles and surrounding parenchyma. The insert on panel XI is a 400X image of an area of parenchymal consolidation. The white arrow on panel XII points to an area of perivascular infiltration. The black arrow on panel XII points to an area of peribronchiolar infiltration. **B. Combined pulmonary pathology scores.** Mean combined pulmonary pathology scores of hematoxylin and eosin-stained lung sections were calculated for experimental groups of 5 or 6 rats. Error bars denote SEM. Statistical significance was assessed by Mann-Whittney U test. *p < 0.05, **p < 0.01.

In agreement with the original descriptions of pulmonary pathology for cotton rats infected with HAdV-C5 reported by Prince *et al*. (25), we observed histopathologic changes in two phases, an early phase of monocyte-macrophage and neutrophil infiltration affecting alveoli, bronchiolar epithelium, and peribronchiolar regions spanning days 1-3; and a later phase of lymphocytic infiltration affecting peribronchiolar and perivascular regions apparent at day 6 pi. Contrary to what we were expecting based on the *in vitro* data, the average combined pathology scores were higher at all time points sampled for rats infected with PG II strain NHRC 42606 (Figure 6B). Differences were statistically significant at days 1, 3, and 6. On day 1 pi, rats infected with this strain showed a stronger and more significant increase in interstitial pneumonia (inflammatory cell infiltration and thickening of alveolar walls), and alveolitis (inflammatory cell infiltration within the alveolar spaces) than rats infected with PG I strain CL68578. On day 3 pi there was a significant enhancement in interstitial and alveolar cell infiltration in rats infected with strain NHRC 42606, with some of them exhibiting complete consolidation of the lung parenchyma. This increase in lung pathology correlated with the initiation of mortality at 3 dpi recorded in 2 animals infected with this virus.

Rats euthanized on day 6 presented the maximum overall pathology for both infected groups. There was evidence of progression towards a marked increase in peribronchiolitis and perivasculitis (particularly in the PG II strain NHRC 42606-infected rats), and a partial resolution of the parenchymal infiltration (milder interstitial pneumonia and alveolitis).

The levels of pulmonary proinflammatory cytokine/chemokine gene expression were determined by qPCR in lung homogenate supernatants from animals euthanized at 1, 2, 3, and 6 dpi. Statistically significant differences between groups were only detectable at 1 dpi when higher levels of IL-6, IFNβ, MCP-1, IP-10, and RANTES (CCL5) transcripts were observed in samples from PG II strain NHRC 42606-infected animals. Although the mean fold increase values for IFNγ, TNFα, and GRO were also higher in the PG II-infected cotton rats, differences between groups were not statistically significant (Figure 7). These results are consistent with the more pronounced cellular infiltration observed in the lungs of rats infected with the PG II virus. Cytokine transcript levels at the other time points post infection were not statistically different between groups (data not shown).

**Figure 7:**
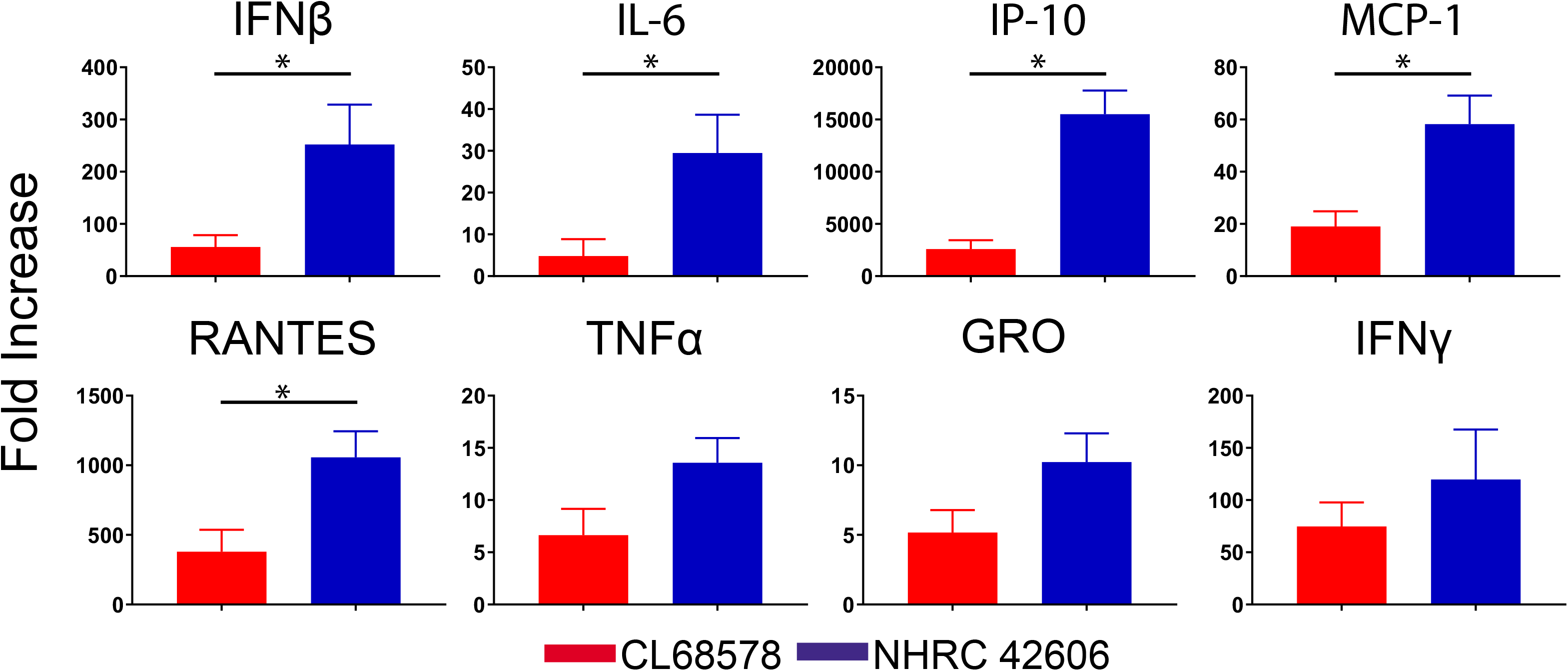
Proinflammatory cytokine and chemokine gene expression in infected cotton rat lungs at 1 dpi. The levels of gene expression in lung homogenates of cotton rats infected with the PG I strain CL68578, PG II strain NHRC 42606, or mock-infected were determined by RT-qPCR. n= 5 or 6 rats per group. Error bars denote SEM. Statistical significance was assessed by Student’s *t* test. p < 0.05.

## DISCUSSION

Evidence of intratypic genetic variably for HAdV-E4 has been reported in the literature since the 1980s, with the description of multiple genomic variants identified by RFLP analysis of serologically indistinguishable strains isolated from cases of respiratory illness and conjunctivitis. However, the clinical significance of this naturally occurring variation has remained elusive for this and other pathogenic HAdV types. In this paper, deeper sequence analysis and proteotyping of a panel of 47 HAdV-E4 whole genome sequences representing 45 strains enabled us to demonstrate additional differences between the two major phylogroups (PG) of HAdV-E4 described by Gonzalez *et al*. (14), and also among the examined genomic variants clustering within each of the two evolutionary lineages.

While the genomes of strains clustering into PG I and PG II are overall 94.5% identical, we previously identified sequence differences between the two lineages mapping throughout the entire genome. These included numerous single nucleotide polymorphisms, insertions and deletions in regions of the genome that have the potential to contribute to *in vivo* phenotypes relevant to pathogenesis (14, 18, 26–29).

Our proteotyping analyses of selected coding regions confirmed that PG I and PG II encode different variants not only for proteins involved in viral replication (E1A, E1B-55K, DNA Pol, L4-100K, and E4-34K), but also for proteins with immunomodulatory functions such as E1B-19K (30) and the E3-encoded gp19K, 14.7K, and CR1β (31–34). Using a subset of previously sequenced strains that represent different genomic variants in the two phylogroups, we conducted a comparative evaluation of *in vitro* and *in vivo* phenotypes reflecting replicative fitness and pathogenicity. We demonstrated marked differences between the two genetic lineages of HAdV-E4.

Compared to many RNA viruses for which genetic determinants of fitness and virulence have been identified and links between genetic variability and phenotypic variability have been demonstrated (35–39), relatively little is known about the implications of genetic variability for DNA viruses of medical importance. Recent applications of genomics approaches to the study of various aspects of human herpesvirus pathogenesis have advanced the understanding of the magnitude and implications of genetic diversity for members of this group (40–44). In addition, important studies have identified gene variations of myxoma virus field strains that may be responsible for changes in virulence (45). To our knowledge, this paper is the first to report phenotypic correlates of naturally occurring intratypic genetic variability for human adenoviruses associated with disease. The work presented here demonstrates distinct *in vitro* and *in vivo* phenotypes relevant to pathogenesis for HAdV-E4 strains belonging to PG I and PG II. We show that in cell culture, PG I strains are fitter viruses than PG II strains, exhibiting larger plaque sizes, enhanced lateral spread, faster plaque development and replication, enhanced cell killing, and earlier release of infectious viral progeny. The documented coding variations in the E1, E2, and E4 regions among other shown in Table 1 segregate with these phenotypic differences. A few of these regions have been previously identified experimentally to contribute to replicative fitness. Through the generation of compensatory mutants with a restored large plaque phenotype compared to the parental ADP-deficient mutant of HAdV-C5 *dl327*, Subramanian *et al*. identified E1A, E1B19K, E1B55K, and E4Orf3 as genes influencing the spread phenotype (46). Importantly, these authors also identified compensatory mutations in the penton base and fiber genes that warrant further investigation for HAdV-E4 strains.

As documented by our group and others, genomes of HAdV-E4 strains in PG I and PG II also differ in the sequences of their respective inverted terminal repeats (ITRs) (14, 26, 27). Wunderlich *et al*. reported an increased genome replication rate for a HAdV-B35-based vaccine vector carrying the alternative ITR sequence 5’-CTATCTAT-3’ (47). Interestingly, they also described the presence of this alternative ITR sequence in the genomes of several HAdV types, including HAdV-E4 PG I strains RI-67 and CL68578, which in our *in vitro* experiments had faster replication rates and enhanced lateral spread compared to the PG II viruses.

The deeper examination of strains with apparent intermediate phenotypes for lateral spread and plaque size (RI-67) or cell killing (NY-15-4054) as well as of strains encoding different protein variants in loci of interest will facilitate the finer mapping of genetic determinants of replicative fitness.

In the cotton rat model of HAdV respiratory infection the pathological response to HAdV-E4 resembled that previously described for HAdV-C5 (25). Importantly, the model allowed us to observe marked differences in induced pulmonary pathology between the examined PG I and PG II viruses. This demonstrates the value of the cotton rat as a permissive host for experimental *in vivo* examination of the effects of genetic variability of HAdV-E4 on virulence and for the identification of candidate genetic determinants of pathogenicity for this unique HAdV type.

Surprisingly, the differences in replicative fitness documented *in vitro* between PGs did not correlate with the differences in virulence observed in the cotton rat model. These data strongly implicate a differential involvement of the host immune response in the pathogenesis of the documented pulmonary disease rather than a direct viral cytopathic effect. The detection of significant differences in the levels of gene expression for proinflammatory cytokines and of chemokines that regulate the migration and infiltration of monocytes, macrophages, and T cells (MCP-1*/*CCL2, IP-10, and RANTES/CCL5) are consistent with the observation of enhanced pulmonary pathology in rats infected with the PG II virus.

With the exception of RIDα, all of the E3 proteins encoded in the genomes of CL68578 (PG I) and NHRC 42606 (PG II) are different, i.e. they represent different proteotypes, so it is likely that the two examined viruses differ in their ability to elicit and modulate host responses to infection.

There is a documented higher representation of PG II in HAdV-E4 positive clinical specimens obtained from symptomatic patients (11, 14, 16–18) which suggests a higher “epidemiologic fitness” and/or “transmission fitness” (48) for this clade. PG II strain NHRC 42606 was more virulent than PG I strain CL68578 in our *in vivo* experiments, as evidenced by the more exacerbated inflammatory response in the lung of infected animals and associated fatalities. Interestingly, the studies conducted by Russell *et al*. during 2004 in a cohort of unvaccinated military recruits in basic training at the Marine Corps Recruit Training Command, San Diego, California revealed the efficient transmission of PG II genomic variant 4a (represented in our in vitro studies by strain NHRC 22652) in that unique setting, and collected data that implicated the presence of infectious virus in various fomites and environmental samples in the process (49).

Further work using isogenic virus mutants carrying some of the identified mutations that encode different proteotypes in various regions of the viral genome will be necessary to dissect the contribution of individual coding regions to the differences in observed pulmonary pathology phenotypes between PG I and PG II viruses. Of particular interest are the variants encoded in regions implicated in the modulation of host responses to infection, such as E1, L4, E3, and VA RNA (50–55).

Using the Syrian hamster model of HAdV respiratory infection, Radke *et al*. reported enhanced pulmonary pathology for a naturally occurring genomic variant of HAdV-B14, 14p1, whose genomic sequence differs from that of the prototype strain de Wit in regions of the genome including E1 B and whose distinct *in vitro* phenotypes include a decreased expression of E1B-19K, the viral Bcl2 homolog, and the formation of larger plaques (56).

Like the mouse, the cotton rat has been shown to be a powerful host to evaluate the effect of early genes in HAdV-C5 pathogenesis (57–60). Our experiments demonstrate that *Sigmodon hispidus* is permissive to HAdV-E4 infection and we expect the model to be particularly useful for examination of the role of the species HAdV-E-specific E3-CR1 proteins that are predominantly expressed from late transcripts (61,62). Importantly, the biological function of E3CR1δ/E3-30K has not been identified to the present (61). The work of Li and Wold (59) and preliminary studies in our laboratory (Kajon, personal communication) indicate that HAdV-E4 does not encode an ADP homolog.

Our studies uncover a powerful experimental system and provide a strong foundation for further exploration of specific genotype-phenotype associations for HAdV-E4 and possibly for HAdVs in general. Our various low passage clinical isolates of HAdV-E4 and their corresponding whole genome sequence provide a unique resource to examine other phenotypes both *in vitro* and *in vivo* to define system-specific correlations of fitness and virulence.

The identification of genetic determinants of fitness and virulence will be extremely useful for the design of HAdV-E4-based vectors and for the genetic manipulation of viruses for attenuation and use in live vaccine formulations. In addition, the identification of genetic factors that influence pathogenicity can potentially contribute to the development of molecular typing tools designed to identify mutations associated with increased virulence at loci of special interest.

## MATERIALS AND METHODS

### Genetic analysis and proteotyping

To identify the distribution of nucleotide differences among HAdV-E4 genomes, the complete genomic sequences of HAdV-E4 strains in phylogroups (PG) I and II characterized by Gonzalez *et al*. (14) were aligned using MAFFT (63). Sequences were ordered following the branching pattern of the phylogenetic tree inferred with MEGA7 (64). Nucleotide sites were color-coded per genome in comparison to the consensus inferred from the multiple sequence alignment. The comparisons were performed with the R package APE (https://cran.r-project.org/web/packages/ape/) to identify by position the consensus nucleotide and the mutations. The predicted amino acid sequence divergence for selected coding regions among PG I and PG II HAdV-E4 strains was examined using the method of proteotyping adapted from Obenauer *et al*. (65). The concatenated protein sequences of E1A, E1B 19K and 55K, DNA polymerase, L4 100K, E3 proteins, and E4 34K were analyzed. Amino acid signatures were derived from a phylogenetic tree-guided sequence alignment, indicating amino acid sites with polymorphisms relative to the most frequently occurring residues with a frequency-based color coding.

### Cell Culture

Human A549 type II alveolar epithelial cells (ATCC CL-185) were grown in Minimum Essential Medium (MEM, Gibco) supplemented with 8% heat-inactivated newborn calf serum (HI-NBCS, Rocky Mountain Biologicals), 1.5 g/ml Na2CO3 (Amresco), 2 mM L-glutamine (Corning), 10 U/ml penicillin (Corning), 10 μg/ml streptomycin (Corning), and 25 mM HEPES (Corning). Human Calu-3 airway epithelial cells (ATCC HTB-55) were grown in RPMI-1640 medium (Corning) supplemented with 10% fetal bovine serum (FBS, Atlanta Biologicals), 2 mM L-glutamine, 10 U/ml penicillin, 10 μg/ml streptomycin, and 25 mM HEPES. Cotton rat lung epithelial cells (CRLC, ATCC PTA-3930) were grown in MEM (Gibco) supplemented as above except with 10% (v/v) heat-inactivated FBS (Atlanta Biologicals) instead of HI-NBCS. For virus infections, all cell lines were cultured in the appropriate medium with the serum content reduced to 2%. The cell lines were confirmed to be Mycoplasma sp. free by testing using the ABM Mycoplasma PCR Detection Kit (G238, Applied Biological Materials Inc., Canada).

### Viruses

Low passage virus stocks of clinical isolates were produced in A549 cells. After 3 rounds of freeze/thawing, cell debris was removed by extraction with of H2O-saturated chloroform. After venting overnight at room temperature to remove traces of chloroform, infectious titers were determined by standard plaque assay on A549 cell monolayers in 6-well plates. The prototype strain RI-67 of PG I was obtained from the collection of Goran Wadell, Umeå University, Sweden with an unknown history of passage. The PG I 4p vaccine strain CL68578 was derived from the pVQ WT #11183 genomic clone obtained from ABL, Inc. Briefly, the genomic clone was released from the bacterial plasmid by digestion with Pac I (New England Biolabs), and 1 μg of linearized DNA was transfected into A549 cells using Effectene Transfection Reagent (Qiagen) following the manufacturer’s recommendations. The resulting infectious virus was passaged minimally in A549 cells for stock production.

#### Cell-based assays

##### Plaque assay

Confluent A549 cell monolayers in 6 well plates were inoculated with 100 μl of serially diluted virus (1:10 in phosphate buffered saline (PBS). After adsorption for 1 hour at 37°C and 5% CO_2_ with periodic rocking, monolayers were overlaid with 2 ml of a solution consisting of equal parts of 1.4% (w/v) Low Melt Agarose (G-BioSciences) and 2X MEM (Gibco) supplemented with 4% (v/v) HI-NBCS, 4.0 mM L-glutamine, 20 U/ml penicillin, 20 μg/ml streptomycin, and 25 mM MgCl2. At 7 dpi, infected monolayers were fixed with 1% formaldehyde (Sigma)/ 0.15 M NaCl and stained with crystal violet (EMD-Millipore Sigma).

##### Plaque development assay

A549 cells were seeded in 15 mm x 60 mm dishes to be confluent on the day of infection. Stocks were diluted in PBS so that approximately equal number of PFUs of each strain were used to inoculate each dish in biological triplicate. Growth medium was aspirated, and the cells were inoculated with diluted virus and synchronized on ice for 10 minutes. Inoculated cells were then incubated for two hours at 37°C 5% CO_2_. Following absorption, 5 ml of overlay medium (described above) were added and allowed to solidify at room temperature, then incubated at 37°C and 5% CO_2_. At 3 dpi, an additional overlay containing 0.0015% neutral red dye (JT Baker Chemical Co.) was added. Starting a 4 dpi, plaques were counted daily using a lighted background and marked with a pen. The assay was terminated when no more plaques developed. The data are presented as percentage of the final titer (total number of plaques observed on a given day divided by the total number of plaques observed on the final day of the assay) on each day (21).

##### Plaque size assays

A549 cells in 6-well plates were infected with approximately 20-40 PFU of each virus strain in a volume of 100 μl. After adsorption for 1 hour at 37°C with occasional rocking, the infected monolayers were overlaid as described above. At 7 dpi, prior to fixation, individual plaques were photographed using an Olympus CKX41 microscope equipped with an Infinity 2-3 digital camera (Teledyne Lumenera) and a 4X objective. Images were captured using the Infinity Capture software V6.5.4 (Teledyne Lumenera). The plaque assays were then fixed, stained with crystal violet, and photographed using a copy stand and a transilluminator with the camera in a fixed position. The average plaque size in arbitrary units was determined using the FIJI image analysis software (66).

##### Growth kinetics studies

A549 cells seeded in 10 mm x 35 mm cell culture dishes at 80% confluence were inoculated with HAdV-E4 strains at a MOI of 1 PFU/cell in biological triplicate, incubated on ice for 10 minutes, then absorbed for one hour at 37°C and 5% CO2 with periodic rocking. Unbound virus was removed by aspiration, and the infected cell monolayers were washed twice with PBS. Cells were then replenished with A549 Infection Medium (MEM + 2% HI-NBCI + supplements) and incubated at 37°C with 5% CO2. At various times post infection, infected cell monolayers were frozen at −80°C and subjected to three freeze/thaw cycles. Cell debris was clarified by centrifugation at 300 x g for 10 minutes, and infectious virus titers were determined by plaque assay as described above.

CRLC monolayers were infected with strains CL68578 or NHRC 42606 at a MOI of 5 PFU/cell in biological triplicate, incubated on ice for 10 minutes, then absorbed for one hour at 37°C and 5% CO2 with periodic rocking. At 0, 12, 24, 48 and 72 hpi, cell monolayers were processed as described above for titration of infectious virus yields.

##### Virus spread assays

A549 cells seeded on 48 well plates to be 100% confluent at the time of infection were inoculated with 100 μl of virus stock diluted in PBS for MOIs of 10, 1, 0.1, 0.01, and 0.001 PFU/cell. After adsorption for 1 hour at 37°C, the inoculum was removed by aspiration, cell monolayers were washed twice with PBS, replenished with A549 Infection Medium, and incubated at 37°C and 5% CO2. At 7 dpi cell monolayers were fixed with 1% formaldehyde in 0.15 M NaCl, and stained with crystal violet.

##### Cell viability assays

Intracellular ATP levels were measured at various times post infection as indicators of the presence of metabolically active cells and thus of cell viability (67) using the luminescent cell viability CellTiter-Glo 2.0 assay (Promega) according to the manufacturer’s specifications. Briefly, A549 cells were seeded in 96 well dishes to be 75% confluent on the day of infection. Cells were inoculated at a MOI of 10 PFU/cell with virus diluted in PBS or mock-infected with PBS in biological quadruplicate. After adsorption for 2 hours at 37°C the inoculum was removed, monolayers were washed twice with PBS, and then then replenished with 50 μl of A549 Infection Medium. Relative luminescence was measured in a Luminoskan Ascent Type 392 (ThermoFisher Scientific). Data are reported as mean relative luminescence units ± SEM.

To further characterize the cell killing phenotype of PG I and PG II strains, LDH release from lysed cells was assessed using the colorimetric CytoTox 96 Non-radioactive Cytotoxicity Assay Kit (Promega). Briefly, A549 cells seeded in 48-well plates to be confluent at the time of infection were inoculated with virus diluted in PBS at a MOI of 1 PFU/cell in biological triplicates. After adsorption for 1 hour at 37°C the inoculum was removed, monolayers were washed twice with PBS and then replenished with infection medium. At 1, 2, and 3 dpi, the supernatant was removed and centrifuged at 300 x g for 10 minutes. LDH activity was then assessed according to the manufacturer’s directions in technical quadruplicate. Data are reported as mean absorbance at 490 nm ± the standard error of the mean (SEM).

##### Viral progeny release assays in polarized Calu-3 cells

Calu-3 cells were seeded onto transwell inserts (0.4 μm pore size, Corning) at a density of 5 x10^5^ cells/well, and Calu-3 growth medium was added to both the apical and basal compartments. Two days post seeding, the cells were washed with PBS++ (Dulbecco’s Phosphate-Buffered Saline with Magnesium and Calcium, Gibco). All liquid was then removed from the apical chamber, and the cells were grown at an air/liquid interphase for 10 days with medium changes every 48 hours. Polarization of Calu-3 cells was evaluated by measuring the transepithelial resistance (TER) using a Millicell ERS meter (Millipore). Cells were considered polarized when TER values reached 500-600 Ωcm^-2^.

For infection, medium was removed from the basal compartment, and Calu-3 infection medium was added. Virus diluted in PBS++ for a MOI of 10 PFU/cell was then added to the apical compartment. After absorption at 37°C, 5% CO2 for 1 hour, the inoculum was removed, and the cells were washed with PBS++. All liquid in the apical and basal compartments was removed, and fresh Calu-3 infection medium was added to the basal compartment. The cells were then incubated at 37°C, 5% CO2. At 1-3 dpi, the infected cells were processed by washing the apical compartment twice with a total of 400 μl of PBS ++. Infectious virus titers in the apical wash were determined by plaque assay.

#### Cotton rat studies

All animal studies were conducted under applicable laws and guidelines and after protocol approval by the Sigmovir Biosystems, Inc. (SBI) Institutional Animal Care and Use Committee. Cotton rats (*Sigmodon hispidus*) obtained from SBI’s inbred colony were housed in large polycarbonate cages and fed a diet of standard rodent chow and water *ad libitum*. The number of animals per experimental group required to observe differences in pulmonary pathology scores was determined to be 6 by power analysis using G*power software v3.1.9.2 (68), The calculations were carried out assuming a significance level of 0.05, 80% power, a signal/noise ratio of 1.5 pathology score units, and a standard deviation of 1.

Four-week-old female cotton rats were inoculated with 5×10^6^ PFUs of PG I strain CL68578 or PG II strain NHRC 42606 in each nare under isoflurane anesthesia. Control mock-infected animals were inoculated in the same manner with chloroform-extracted supernatants of uninfected A549 cells. Cotton rats were sacrificed at 1, 2, 3, and 6 days post infection by CO2 asphyxiation. Lung tissue was collected for downstream analyses as follows: left lobe for assessment of viral load; left lingular lobe for mRNA reverse transcription for cytokine gene expression; and right lung for histopathology. All lung lobes except for the right lobe were snap-frozen in liquid nitrogen and stored at −80°C until processed. The right lung was inflated with 10% neutral buffered formalin.

### Assessment of pulmonary viral load

Frozen left lobe sections were weighed and then homogenized in 1mL of A549 infection medium using a TissueLyser II apparatus (Qiagen) and 3 cycles of 30 hz for 30 seconds, chilling on ice between each cycle. Homogenates were centrifuged at 16,000 X g for 10 minutes, and the supernatant was aliquoted into three separate tubes of equal volume.

One aliquot of lung homogenate supernatant was processed for assessment of infectious viral load by plaque assay. Supernatants were serially diluted ten-fold in PBS and 100 μl were plated in triplicate onto monolayers of A549 cells in 6-well plates. The assays were carried out as described above.

At DiaSorin Molecular (Cypress, CA), DNA was extracted from 200 μl of another aliquot of lung homogenate supernatant using the MagNA Pure Total Nucleic Acid Isolation protocol and reagents on a MagNA Pure LC 2.0 instrument (Roche Diagnostics). Prior to extraction, 5 μl of the Simplexa Extraction and Amplification control DNA (SEAC, DiaSorin Molecular) were added to each sample. Samples were eluted in 50 μl.

The real-time qPCR assay was carried on the LIAISON^®^ MDX instrument (DiaSorin Molecular) using the 96 well Universal Disc (DiaSorin Molecular) and DiaSorin Molecular Adenovirus analyte-specific reagents. The adenovirus 3’ Hexon primer pair and the adenovirus 5’ Hexon primer pair with FAM-labeled forward primers were used to amplify and detect conserved regions at the 3’ and 5’ ends of the HAdV hexon gene. A Control SEAC Primer Pair with a Quasar^®^ 760-labeled forward primer was used to amplify and detect the SEAC DNA fragment. The 5’ and 3’ hexon primer pairs are a proprietary bi-functional fluorescent primer-probe set targeting the 3’ and 5’ regions of the HAdV hexon gene. Each reaction contained 4 μl of 2.5X Universal Master Mix, 0.2 μl of Adenovirus 3’ Hexon primer pair, 0.2 μl of Adenovirus 5’ Hexon primer pair, 0.2 μl of internal control primer, 0.4 μl of water, and 5 μl of extracted template, in a total reaction volume of 10 μl. PCR amplification with real-time detection was performed using the following cycling conditions: denaturation at 97°C for 120 seconds (1 cycle); denaturation at 97°C for 10 seconds followed by annealing/extension at 60°C for 30 seconds (40 cycles). Fluorescence was measured at the end of each annealing/extension cycle with quantitative values calculated by the LIAISON^®^ MDS software.

Standard curves were generated using highly pure genomic DNA extracted from either PG I vaccine strain CL68578- or PG II clinical isolate NHRC 42606 −infected A549 cells as previously described (69). The calibrator values ranged from 1.2×10^6^ to 1.2×10^2^ genome copies (cp) /ml for CL68578 and from 2.8×10^6^ to 2.8×10^2^ cp/ml for NHRC 42606. Each standard was extracted in singlicate and each extract amplified in quadruplicate in a single run. Quantitative analysis was performed by the LIAISON^®^ MDX Software. Standard curves were used for all viral load determinations. Viral loads were expressed as genome copies per gram of tissue.

### RNA Isolation and Reverse Transcription

Total RNA was extracted from the left lingular lung lobes using the RNAqueous kit (ThermoFisher Scientific). Frozen lung sections were homogenized in 1 ml of Lysis/Binding solution in a TissueLyser II (Qiagen) with three cycles of at 30 Hz for 30 seconds, with chilling on ice between cycles. Samples were centrifuged at 16,000 x g for 2 minutes at 4°C and the supernatant was used for total RNA isolation according to the manufacturer’s recommendations. Residual DNA was removed from the extracted RNA with the TurboDNA-free kit (ThermoFisher Scientific). The removal of contaminating DNA was verified by PCR amplification of the rig/S15 gene using the forward (5’ – 3’) primer TTC CGC AAG TTC ACC TAC C and the reverse (3’ – 5’) primer CGG GCC GGC CAT GCT TTA CG with GoTaq Flexi DNA polymerase (Promega) and the following cycling conditions: 1 cycle of 95°C, 5 minutes; 50 cycles of 95°C for 30 seconds, followed by 57°C for 30 seconds, then 72°C for 30 seconds; 1 cycle of 72°C for 5 minutes. Reverse transcription was then performed on 1 μg of DNAse-treated RNA using the RetroScript Reverse Transcription kit (Invitrogen/ThermoFisher Scientific) and oligo(dT) primers following the manufacturer’s recommendations.

### Histological analysis

Dissected right lung lobes were inflated with 10% neutral buffered formalin and immersed in formalin for fixation. Tissues were then shipped to Histoserv, Inc. (Germantown, MD) for paraffin-embedding, sectioning, and staining with hematoxylin and eosin. The stained lung sections were scored blindly by co-author J.C.G.B. on a scale of 0 to 4 (absent, minimal, mild, moderate, and marked) for the following indices of pulmonary inflammation: peribronchiolitis (peribronchiolar inflammatory cell infiltration), perivasculitis (inflammatory cell infiltration around the small blood vessels), interstitial pneumonia (inflammatory cell infiltration and thickening of alveolar walls), and alveolitis (inflammatory cell infiltration of alveolar spaces). The average combined pathology score was calculated from these four individual components.

### Immunohistochemistry

Services were contracted from the Veterinary Diagnostic Laboratory, University of Minnesota (https://www.vdl.umn.edu/) where paraffin-embedded lung sections on glass slides were processed for staining with AdV Mab MAB805 (Millipore) following an in-house protocol. Images from stained sections were acquired in a Nikon Eclipse E600 microscope equipped with a Nikon DXM1200F digital camera at 40X magnification.

### Analysis of cytokine/chemokine gene expression in lung tissue

cDNA obtained as described above was diluted to 0.1 μg /ml and 3 μl were used as template for each 25 μl qPCR reaction using primers and cycling conditions described previously for IL-6, IFNβ, IFNγ, TNFα, MCP-1, IP-10, RANTES, and GRO (70–72). qPCR reactions were carried out using the QuantiFast SYBR Green PCR kit (Qiagen, Valencia, CA) in a MyiQ single color Real-time PCR cycler (Bio-Rad, Temecula, CA). Relative quantification of PCR products was performed by normalizing to β-actin using the ΔΔC_T_ method. Data are presented as mean fold induction ± SEM.

### Statistical Analysis

Unless otherwise noted, all statistical analyses were carried out using GraphPad Prism v8.1.0 for Microsoft Windows (GraphPad, https://www.graphpad.com). p values < 0.05 were considered statistically significant.

## ACKNOWLEDGMENTS

The authors wish to thank Bojana Rodic-Polic, DiaSorin Molecular (Cypress, CA) for the execution of qPCR assays to assess viral load in cotton rat lung homogenates and Xioyan Lu at CDC (Atlanta, GA) for the qPCR assessment of viral load for our infectious stocks. We also especially thank Kathy Spindler and Matt Weitzman for critical reading of the manuscript and constructive input.

C.R.B. was supported by the University of New Mexico Infectious Diseases and Inflammation NIH Training Grant T32-AI007538.

A.E.K. and C.R.B conceived the study. C.R.B., G.G., W.Z, A.K., D.S., J.C.G.B. and A.E.K. performed investigations for this study. C.R.B. and A.E.K. wrote the manuscript, and all authors reviewed and edited the manuscript.

